# Personalized prediction of rehabilitation outcomes in multiple sclerosis: a proof-of-concept using clinical data, digital health metrics, and machine learning

**DOI:** 10.1101/2020.03.26.010264

**Authors:** Christoph M. Kanzler, Ilse Lamers, Peter Feys, Roger Gassert, Olivier Lambercy

**Author notes:** **Corresponding author:** Christoph M. Kanzler, Rehabilitation Engineering Laboratory, ETH Zürich, BAA C 307.1, Lengghalde 5, 8008 Zürich, Switzerland.

## Abstract

**Background:** A personalized prediction of upper limb neurorehabilitation outcomes in persons with multiple sclerosis (pwMS) promises to optimize the allocation of therapy and to stratify individuals for resource-demanding clinical trials. Previous research identified predictors on a population level through linear models and clinical data, including conventional assessments describing sensorimotor impairments. The objective of this work was to explore the feasibility of providing an individualized and more accurate prediction of rehabilitation outcomes in pwMS by leveraging non-linear machine learning models, clinical data, and digital health metrics characterizing sensorimotor impairments.

**Methods:** Clinical data and digital health metrics were recorded from eleven pwMS undergoing neurorehabilitation. Machine learning models were trained on data recorded pre-intervention. The dependent variables indicated whether a considerable improvement on the activity level was observed across the intervention or not (binary classification), as defined by the Action Research Arm Test (ARAT), Box and Block Test (BBT), or Nine Hole Peg Test (NHPT).

**Results:** In a cross-validation, considerable improvements in ARAT or BBT could be accurately predicted (94% balanced accuracy) by only relying on patient master data. Considerable improvements in NHPT could be accurately predicted (89% balanced accuracy), but required knowledge about sensorimotor impairments. Assessing these with digital health metrics instead of conventional scales allowed increasing the balanced accuracy by +17% . Non-linear machine-learning models improved the predictive accuracy for the NHPT by +25% compared to linear models.

**Conclusions:** This work demonstrates the feasibility of a personalized prediction of upper limb neurorehabilitation outcomes in pwMS using multi-modal data collected before neurorehabilitation and machine learning. Information from digital health metrics about sensorimotor impairment was necessary to predict changes in dexterous hand control, thereby underlining their potential to provide a more sensitive and fine-grained assessment than conventional scales. Non-linear models outperformed ones, suggesting that the commonly assumed linearity of neurorehabilitation is oversimplified.

clinicaltrials.gov registration number: NCT02688231

## 1 Introduction

Multiple sclerosis (MS) is a chronic neurodegenerative disorder with a prevalence of 2.2 million worldwide [1]. It disrupts a variety of sensorimotor functions and affects the ability to smoothly and precisely articulate complex multi-joint movements, involving for example the arm and hand [2]. This strongly affects the ability to perform daily life activities, leads to increased dependence on caregivers, and ultimately a reduced quality of life [3]. Inter-disciplinary neurorehabilitation approaches combining, for example, physiotherapy and occupational therapy have shown promise to reduce upper limb disability [4, 5, 6]. This is reflected by a reduction in sensorimotor impairments and an increase in the spectrum of executable activities, as defined by the International Classification of Functioning, Disability, and Health (ICF) [7].

One of the active ingredients to ensure successful neurorehabilitation is a careful adaptation of the therapy regimen to the characteristics and deficits of an individual (i.e., personalized therapy) [5, 8, 6]. For this purpose, predicting whether a patient is susceptible to positively respond to a specific neurorehabilitation intervention is of primary interest to researchers and clinicians, as it can help to set more realistic therapy goals, optimize therapy time, and reduce costs related to unsuccessful interventions [9, 10, 11, 12]. In addition, it promises to define homogenous and responsive groups for large-scale and resource-intensive clinical trials.

Unfortunately, knowledge about predictors determining the response to neurorehabilitation is limited in pwMS [13, 14, 15]. So far, most of the approaches focused on establishing correlations between clinical variables at admission and discharge on a population level. This allowed the identification of, for example, typical routinely collected data (e.g., chronicity) and the severity of initial sensorimotor impairments as factors determining the efficacy of neurorehabilitation [5, 13, 14, 15]. However, identifying trends on a population level has limited relevance to actually inform daily clinical decision making.

Predicting therapy outcomes at an individual level promises to provide clinically-relevant information [9, 10, 11, 12], but requires appropriate modeling and evaluation strategies that go beyond the commonly applied linear correlation analyses. More advanced approaches are necessary to account for potentially non-linear relationships and the high behavioral inter-subject variability commonly observed in neurological disorders. In addition, the severity of sensorimotor impairments is often not characterized in a sensitive manner, which might limit their predictive potential. This stems from assessments of sensorimotor impairments commonly applied in clinical research (referred to as conventional scales) providing only coarse information, as they usually rely on subjective evaluation or purely timed-based outcomes that are not able to describe behavioural variability [16, 17].

Machine learning allows an accurate and data-driven modeling of complex non-linear relationships, which offers high potential for a precise and personalized prediction of rehabilitation outcomes [18, 19]. Similarly, digital health metrics of sensorimotor impairments allow answering certain limitations of conventional scales by providing objective and fine-grained information without ceiling effects [20]. Such kinematic and kinetic metrics have found first pioneering applications in pwMS, allowing to better disentangle the mechanisms underlying sensorimotor impairments [21, 22, 23, 24, 25, 26, 27, 28]. So far, neither of these techniques has been applied for a personalized prediction of rehabilitation outcomes in pwMS.

The objective of this work was to explore the feasibility of predicting upper limb rehabilitation outcomes in individual pwMS by combining clinical data, digital health metrics, and machine learning (Figure 1). For this purpose, clinical data including routinely collected information (e.g., age and chronicity) and conventional assessments were recorded pre- and post-intervention in 11 pwMS that participated in a clinical study on task-oriented upper limb rehabilitation [6]. In addition, digital health metrics describing upper limb movement and grip force patterns were recorded using the Virtual Peg Insertion Test (VPIT), a previously validated technology-aided assessment of upper limb sensorimotor impairments in neurological subjects relying on a haptic end-effector and a virtual goal-directed object manipulation task [24, 29, 28].

**Figure 1:**
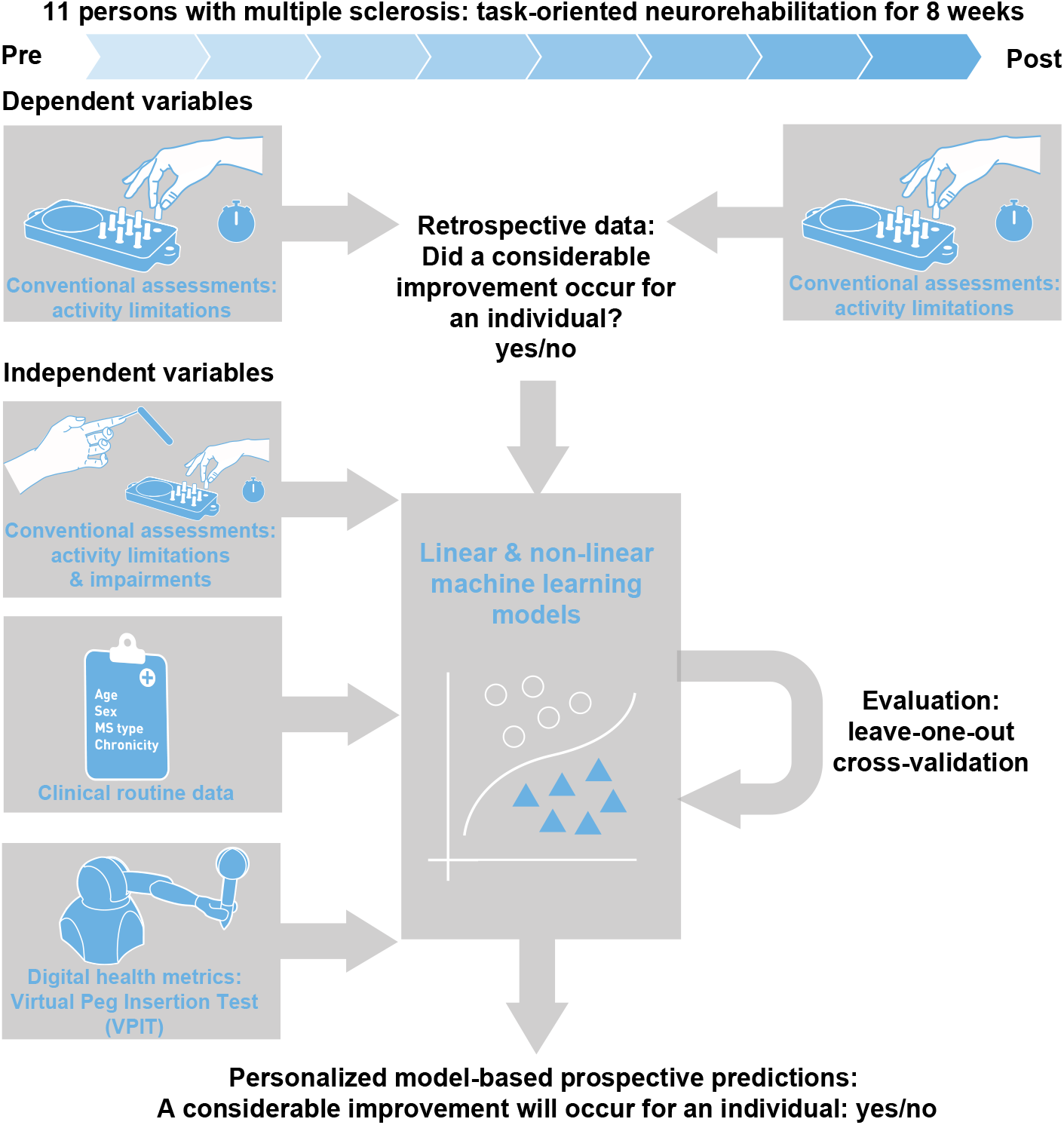
pproach for prediction of neurorehabilitaiton outcomes in persons with multiple sclerosis. Eleven persons with multiple sclerosis were assessed before and after eight weeks of neurorehabilitation. Multiple linear and non-linear machine learning models were trained on different feature sets with data collected before the intervention. This included information from conventional clinical assessments about activity limitations and impairments, clinical routine data, and digital health metrics collected with the Virtual Peg Insertion Test (VPIT). The dependent variable of the models defined whether a considerable improvement in activity limitations occured across the intervention or not. The quality and generalizability of the models was evaluated in a leave-one-subject-out cross-validation.

We hypothesized that 1) machine learning models trained on multi-modal data recorded pre-intervention could inform on the possibility to yield a considerable reduction in upper limb disability due to a specific rehabilitation intervention. Further, we assumed that 2) non-linear machine learning models enable more accurate predictions of rehabilitation outcomes than the more commonly applied linear regression approaches. Lastly, we expected 3) digital health metrics of sensorimotor impairments to provide predictive information that goes beyond the knowledge gained from conventional assessments. Addressing these objectives will pave the way for optimizing neurorehabilitation planning in pwMS and provide further evidence on its efficacy for healthcare practitioners.

## 2 Methods

### Participants

The data used in this work was collected in the context of a clinical study in which the VPIT was integrated as a secondary outcome measure [6]. The study was a pilot randomized controlled trial on the intensity-dependent effects of technology-aided task-oriented upper limb training at the Rehabilitation and MS centre Pelt (Pelt, Belgium). For this purpose, participants were block randomized based on their disability level into three groups receiving either robot-assisted task-oriented training at 50% or 100% of their maximal possible intensity as defined by their maximum possible number of repetitions of a goal-directed task, or alternatively conventional occupational therapy. The training lasted over a period of 8 weeks with 5h of therapy per week. In addition, participants received standard physical therapy focusing on gait and balance. In total, 11 pwMS that successfully performed the VPIT before the intervention were included in the present work. Other study participants did not complete the VPIT protocol due to severe upper limb disability or strong cognitive deficits. Exclusion criteria and details about the study procedures can be found in previous work describing the clinical outcome of the trial [6]. The study was registered at clinicaltrials.gov (NCT02688231) and approved by the responsible Ethical Committees (University of Leuven, Hasselt University, and Mariaziekenhuis Noord-Limburg).

### Conventional assessments

A battery of established conventional assessments was performed to capture the effects of the interventions. On the ICF body function & structure level, impaired sensation in index finger and thumb were tested using Semmes-Weinstein monofilaments (Smith & Nephew Inc., Germantown, USA) [30]. The results of both tests were combined into a single score (2: normal sensation; 12: maximally impaired sensation). Weakness when performing shoulder abduction, elbow flexion, and pinch grip was rated using the Motricity Index, leading to a single score for all three movements (0: no movement; 100: normal power) [31]. The severity of intention tremor and dysmetria was rated during a finger to nose task using Fahn’s Tremor Rating Scale and summed up to a single score describing tremor intensity (0: no tremor; 6: maximum tremor) [32]. Fatigability was evaluated using the Static Fatigue Index that describes the decline of strength during a 30s handgrip strength test (0: minimal fatigability; 100: maximal fatigability; [33]). Cognitive impairment was described using the Symbol Digit Modality Test, which defines the number of correct responses in 90s when learning and recalling the associations between certain symbols and digits [34]. Lastly, the Expanded Disability Status Scale (EDSS; 0: neurologically intact; 10: death) was recorded as an overall disability measure [35].

On the ICF activity level, the Action Research Arm Test (ARAT) evaluated the ability to perform tasks requiring the coordination of arm and hand movements and consists of four parts focusing especially on grasping, gripping, pinching, and gross movements (0: none of the tasks could be completed; 57: all tasks successfully completed without difficulty) [36]. Further, the ability to perform fine dexterous manipulations were described with the time to complete the Nine Hole Peg Test (NHPT) [37, 38]. The capability to execute gross movements was defined through the Box and Block Test (BBT), which defines the number of blocks that can be transferred from one box into another within one minute [39, 36]. The outcomes of the NHPT and the BBT were defined as z-scores based on normative data to account for the influence of sex, age, and the tested body side [39, 40].

### Digital health metrics describing upper limb movement and grip force patterns with the Virtual Peg Insertion Test (VPIT)

The VPIT is a technology-aided assessment that consists of a 3D goal-directed manipulation task. It requires cognitive function as well as the coordination of arm movements, hand movements, and power grip force to insert nine virtual pegs into nine virtual holes [41, 28]. The task is performed with a commercially available haptic end-effector device (PhantomOmni or GeomagicTouch, 3D Systems, USA), a custom-made handle able to record grasping forces, and a virtual reality environment represented on a personal computer. The end-effector device provides haptic feedback that renders the virtual pegboard and its holes. The VPIT protocol consists of an initial familiarization period with standardized instructions followed by five repetitions of the test (i.e., insertion of all nine pegs five times), which should be performed as fast and accurately as possible.

We previously established a processing framework that transforms the recorded kinematic, kinetic, and haptic data into a validated core set of 10 digital health metrics that were objectively selected based on their clinimetric properties [28]. These were defined by considering test-retest reliability, measurement error, and robustness to learning effects (all in neurologically intact participants), the ability of the metrics to accurately discriminate neurologically intact and impaired participants (discriminant validity), as well as the independence of the metrics from each other. The kinematic metrics were extracted during either the transport phase, which is defined as the gross movement from picking up a peg until inserting the peg and requires the application of a grasping force of at least 2N, or the return phase, which is the gross movement from releasing a peg into a hole until the next peg is picked up and does not require the active control of grasping force. Further, the peg approach and hole approach phases were defined as the precise movements before picking up a peg and releasing it into a hole, respectively.

In the following, the definition and interpretation of the ten core metrics of the VPIT are briefly restated (details in previous and related work [28, 20, 42, 43]). The logarithmic jerk transport, logarithmic jerk return, and spectral arc length return are measures of movement smoothness, which is expected to define the quality of an internal model for movement generation producing appropriately scaled neural commands for the intended movement, and leads to bell-shaped velocity profiles in neurologically intact participants. The jerk-based metrics were calculated by integrating over the third derivative of the position trajectory and by normalizing the outcome with respect to movement length and duration. The spectral arc length was obtained by analyzing the frequency content of the velocity profile. Further, the path length ratio transport and path length ratio return described the efficiency of a movement by comparing the shortest possible distance between start and end of the movement phase and relating this to the actually traveled distance. Additionally, the metric velocity max return describes the maximum speed of the end-effector. To capture the behaviour during precise movements when approaching the peg, the metric jerk peg approach was calculated. Lastly, three metrics were calculated to capture grip force coordination during transport and hole approach. In more detail, the force rate number of peaks transport (i.e., number of peaks in the force rate profile) and the force rate spectral arc length transport described the smoothness of the force rate signal and were expected to describe abnormal oscillations in the modulation of grip force during gross arm movements. Additionally, the force rate spectral arc length metric was calculated during the hole approach phase.

After the calculation of the metrics, the data processing framework includes the modeling and removal of potential confounds including age, sex, whether the test was performed with the dominant hand or not, and stereo vision deficits. Lastly, the metrics are normalized with respect to the performance of 120 neurologically intact participants and additionally to the neurologically impaired subject in the VPIT database that showed worst task performance according to each specific metric. This leads to continuous outcome measures in the unbounded interval]— ∞%, +∞%[, with the value 0% indicating median task performance of the neurologically intact reference population and 100% the worst task performance of the neurologically affected participants.

### Data analysis

In order to predict neurorehabilitation outcomes, several machine learning models of different complexity were trained on different feature sets (independent variables) recorded pre-intervention [44]. Knowledge about whether a participant yielded a considerable reduction of disability on the activity level across the intervention or not was used as dependent variable for the models (i.e., supervised learning of features with binary ground truth) [44]. A considerable reduction in activity limitations was defined by comparing the change of a conventional assessments (ARAT, BBT, or NHPT) to their smallest real difference (SRD) [45]. The SRD defines a range of values for which the assessment cannot distinguish between measurement noise and an actual change in the measured physiological construct. Hence, changes across above the SRD were defined as considerable improvements. Conventional assessments describing activity limitations were used and preferred over a characterization of body functions & structures, as improving the former is more commonly the primary target during neurorehabilitation, and conventional assessments of activity limitations provide more sensitive scales (often continuous time-based) than conventional assessment of body functions (often ordinal) [5]. The SRD for the ARAT, BBT, and NHPT were previously defined for neurological subjects as 5.27 points, 8.11 min^-1^, and 5.32 s respectively [46, 47, 48]. Separate machine learning models were trained for each of these conventional assessments, given that individuals might change only selectively in a subset of these.

The data-driven machine learning models allow generating a transfer function that might be able to associate the value of the independent variables recorded pre-intervention to the rehabilitation outcomes (expected reduction in activity limitations: yes or no). The predictive power of the models was evaluated by comparing the ground truth (i.e., whether a subject significantly reduced its activity limitations across the intervention) with the model estimates for each pwMS via a confusion matrix and the balanced accuracy (i.e., average of sensitivity and specificity). As complex machine learning models can theoretically perfectly fit to any type of multi-dimensional data, the models were tested on data that were not used for its training. For this purpose, leave-one-out crossvalidation was applied to train the model on data from all participants except one. Subsequently, this hold-out data set is used to test the generalizability of the model. This process is repeated until all possible permutations of testing and training set are covered and is expected to provide a performance evaluation of the model that is more generalizable to unseen data (i.e., assessments on new patients). In addition, the models were specifically evaluated on individuals that have considerable activity limitations pre-intervention but do not show a positive response to neurorehabilitation (i.e., unexpected non-responders), as such patients are of high interest from a clinical perspective. As we hypothesized that different predictive factors influence rehabilitation outcomes, multiple feature sets (i.e., combination of multiple independent variables) were defined. In more detail, six basic feature sets containing patient master data (MS type, chronicity, age, sex), intervention group, disability level (EDSS and disability information used for block randomization), sensorimotor impairments assessed with conventional scales (motricity index, static fatigue index, monofilament index, symbol digit modality test, Fahn’s tremor rating scale), sensorimotor impairments assessed with the VPIT (ten digital health metrics), and activity limitations assessed with conventional scales (ARAT, BBT, NHPT). Separate machine learning models were trained on these basic feature sets and selected combinations thereof. All feature sets only contained information collected before the intervention.

Four types of machine learning models were used to enable comparisons between linear and non-linear approaches and to ensure the robustness of the results to the model choice. Decision trees were applied, consisting of multiple nodes that involve the binary testing of a feature based on a threshold, branches that define the outcome of the test (value of feature above or below the threshold), and multiple leaf nodes that indicate a classification label (considerable improvement or not) [49]. The number of nodes, the metric that is tested at each node, and the thresholds are automatically chosen based on a recursive statistical procedure that attempts to minimize the overlap between the distributions of the two classes (considerable improvement or not). This model was chosen due to its simplicity, intuitive interpretability, and high generalizability. In addition, k-nearest neighbor (classification based on normalized Euclidean distances) and random forest (combination of multiple decision trees) models were applied [44]. Finally, a standard linear regression approach was used to establish baseline performance values, given that these are the simplest models and are predominantly used in literature [13, 14, 15]. For this approach, the model outputs were rounded to adhere to the binary classification problem.

## 3 Results

The 11 pwMS (7 female) used for the analysis were of age 56.7±14.8 yrs and had an EDSS of 6.1 ± 1.3, (mean±standard deviation; detailed information in Table 1). Given that all participants except two successfully completed all assessments with both upper limbs, 20 data sets were available for analysis. Six, nine, and six of these data sets showed considerable improvements in the ARAT, BBT, and NHPT, respectively. One participant (ID 02) was an unexpected nonresponder, as he had strong activity limitations (admission: ARAT 44, BBT 20 min^-1^, NHPT 140.27s), but did not make considerable improvements during neurorehabilitation (discharge: ARAT 41, BBT 26 min^-1^, NHPT 216.4s).

**Table 1:** Clinical information on persons with multiple sclerosis. Subject 2 was defined as a unexpected non-responder, as he had the strongest activity limitations at admission, but did not respond positively to neurorehabilitation.

| ID | Time | Side | Age<br>yrs | Sex | MS type | Chronicity<br>yrs | Interv. | EDSS<br>0-10 | ARAT<br>0-57 | BBT<br>1/min | NHPT<br>s |
| --- | --- | --- | --- | --- | --- | --- | --- | --- | --- | --- | --- |
| 01 | Pre | Left | 52 | F | RR | 29 | 1 | 7 | 37 | 22 | 45.25 |
| 01 | Post | Left | 52 | F | RR | 29 | 1 | - | 49 | 33 | 41.14 |
| 01 | Pre | Right | 52 | F | RR | 29 | 1 | 7 | 47 | 31 | 24.75 |
| 01 | Post | Right | 52 | F | RR | 29 | 1 | - | 56 | 47 | 23.43 |
| 02 | Pre | Right | 69 | M | PP | 19 | 1 | 7.5 | 44 | 20 | 140.27 |
| 02 | Post | Right | 69 | M | PP | 19 | 1 | - | 41 | 26 | 216.4 |
| 03 | Pre | Left | 25 | F | RR | 6 | 2 | 6 | 52 | 45 | 29.35 |
| 03 | Post | Left | 25 | F | RR | 6 | 2 | - | 57 | 57 | 23.78 |
| 03 | Pre | Right | 25 | F | RR | 6 | 2 | 6 | 53 | 43 | 29.62 |
| 03 | Post | Right | 25 | F | RR | 6 | 2 | - | 55 | 52 | 23.81 |
| 04 | Pre | Left | 42 | F | RR | 1 | 1 | 4 | 56 | 39 | 27.81 |
| 04 | Post | Left | 42 | F | RR | 1 | 1 | - | 56 | 57 | 23.52 |
| 04 | Pre | Right | 42 | F | RR | 1 | 1 | 4 | 54 | 40 | 20.48 |
| 04 | Post | Right | 42 | F | RR | 1 | 1 | - | 55 | 65 | 20.92 |
| 05 | Pre | Left | 56 | F | SP | 10 | 2 | 7 | 49 | 38 | 33.72 |
| 05 | Post | Left | 56 | F | SP | 10 | 2 | - | 54 | 44 | 35.3 |
| 05 | Pre | Right | 56 | F | SP | 10 | 2 | 7 | 29 | 25 | 89.79 |
| 05 | Post | Right | 56 | F | SP | 10 | 2 | - | 40 | 28 | 70.24 |
| 06 | Pre | Left | 65 | M | SP | 19 | 2 | 8 | 52 | 34 | 39.9 |
| 06 | Post | Left | 65 | M | SP | 19 | 2 | - | 54 | 34 | 44.13 |
| 07 | Pre | Left | 63 | F | RR | 8 | 2 | 4.5 | 57 | 60 | 20.84 |
| 07 | Post | Left | 63 | F | RR | 8 | 2 | - | 57 | 49 | 22.28 |
| 07 | Pre | Right | 63 | F | RR | 8 | 2 | 4.5 | 54 | 47 | 35.04 |
| 07 | Post | Right | 63 | F | RR | 8 | 2 | - | 55 | 45 | 27.3 |
| 08 | Pre | Left | 76 | F | RR | 38 | 1 | 5 | 43 | 42 | 27.01 |
| 08 | Post | Left | 76 | F | RR | 38 | 1 | - | 54 | 56 | 24.61 |
| 08 | Pre | Right | 76 | F | RR | 38 | 1 | 5 | 34 | 43 | 34.46 |
| 08 | Post | Right | 76 | F | RR | 38 | 1 | - | 54 | 49 | 23.46 |
| 09 | Pre | Left | 60 | M | PP | 21 | 1 | 7 | 52 | 44 | 31.48 |
| 09 | Post | Left | 60 | M | PP | 21 | 1 | - | 54 | 49 | 35.66 |
| 09 | Pre | Right | 60 | M | PP | 21 | 1 | 7 | 53 | 51 | 25.29 |
| 09 | Post | Right | 60 | M | PP | 21 | 1 | - | 55 | 48 | 41.93 |
| 10 | Pre | Left | 46 | M | PP | 11 | 2 | 5.5 | 55 | 32 | 30.58 |
| 10 | Post | Left | 46 | M | PP | 11 | 2 | - | 56 | 43 | 39.08 |
| 10 | Pre | Right | 46 | M | PP | 11 | 2 | 5.5 | 56 | 35 | 23.23 |
| 10 | Post | Right | 46 | M | PP | 11 | 2 | - | 56 | 53 | 20.83 |
| 11 | Pre | Left | 70 | F | RR | 37 | 3 | 6 | 53 | 45 | 29.86 |
| 11 | Post | Left | 70 | F | RR | 37 | 3 | - | 55 | 39 | 35.28 |
| 11 | Pre | Right | 70 | F | RR | 37 | 3 | 6 | 45 | 42 | 53.21 |
| 11 | Post | Right | 70 | F | RR | 37 | 3 | - | 52 | 41 | 46.19 |

ID: participant identifier. F: female. M: male. Intervention (interv.) group: task-oriented high intensity (1), task-oriented low intensity (2), control (3). RR: relapse remitting. PP: primary progressive. SP: secondary progressive. EDSS: Expanded Disability Status Scale. NHPT: Nine Hole Peg Test. BBT: Box and Block Test. ARAT: Action Research Arm Test. VPIT: Virtual Peg Insertion Test.

The performance evaluation for machine learning models trained on different feature sets and different conventional scores can be found in Table 2 (decision tree), Table SM3 (linear regression), Table SM4 (k-nearest neighbor), and Table SM5 (random forest). One exemplary decision tree trained on digital health metrics and the NHPT is highlighted in Figure 2.

**Figure 2:**
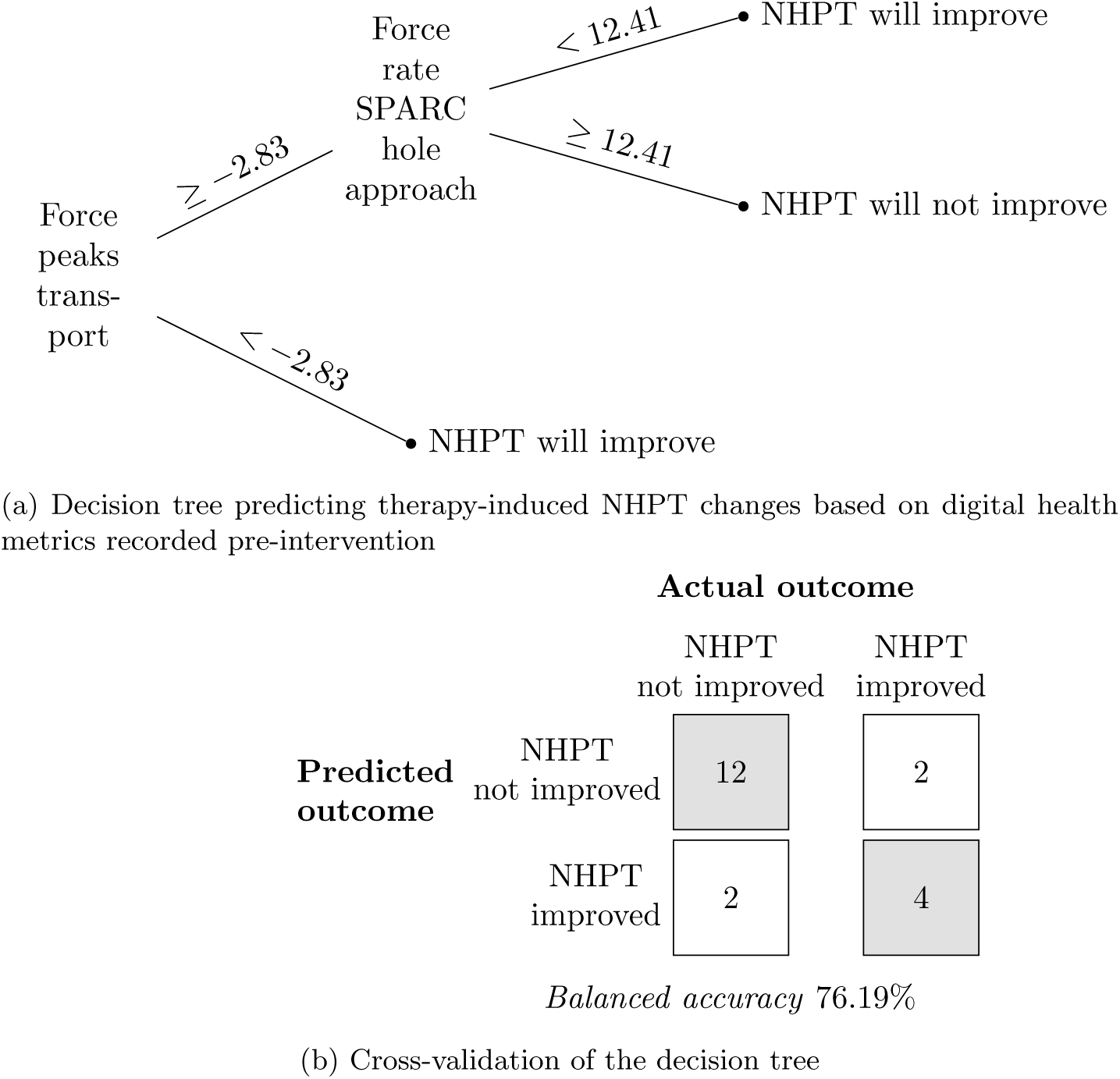
Examplary machine learning model predicting neurorehabilitation outcomes using pre-intervention data. Persons with multiple sclerosis were separated in groups with and without meaningful improvements in activity limitations over an intervention. Multiple machine learning models were trained using different feature sets and independent variables and evaluated using a leave-one-out cross-validation. This exemplary model shows a decision tree (a) that was trained on ten digital health metrics (independent variables) characterizing upper limb sensorimotor impairments, as recorded by the Virtual Peg Insertion Test (VPIT), and the presence or absence of meaningful improvements in activity limitations as observed in the Nine Hole Peg Test (NHPT, dependent variable). Values in the decision tree are reported as normalized VPIT scores. In addition, a confusion matrix (b) and the balanced accuracy (average of sensitivity and specificity) were used within a leave-one-out cross-validation for evaluation purposes. SPARC: spectral arc length.

**Table 2:** Predicting intervention outcomes using pre-intervention data and decision tree models. Multiple machine learning models were trained using different feature sets (independent variables, 1-6). The training label indicated whether a considerable change across intervention was observed in a specific conventional score (dependent variable: ARAT, BBT, or NHPT). The models were evaluated in a leave-one-out cross-validation and specifically tested for one individual with strong activity limitations who did not show improvements across neurorehabilitation (referred to as unexpected non-responder). Feature set nomenclature: 1: patient master data (ms type, chronicity, age, sex). 2: intervention group. 3: disability (EDSS, disability group). 4: Conventional scales of body functions (motricity index, static fatigue index, monofilament index, symbol digit modality test, Fahn’s tremor rating scale). 5: Digital health metrics of sensorimotor impairments (ten VPIT metrics). 6: Conv. scale of activity (ARAT, NHPT, BBT). The best performing (accuracy and unexpected non-responder) models relying on the least amount of features are highlighted in bold for each conventional scale.

| Machine learning: decision tree models |  |  |  |  |  |  |
| --- | --- | --- | --- | --- | --- | --- |
| Feature sets | All participants |  |  | Unexpected non-responder |  |  |
|  | Outcome prediction for |  |  | Outcome prediction for |  |  |
|  | ARAT | BBT | NHPT | ARAT | BBT | NHPT |
|  | Balanced accuracy (%) |  |  | Correct (yes/no) |  |  |
| 1 | **88** | **89** | 33 | y | y | y |
| 2 | 31 | 15 | 33 | y | n | y |
| 3 | 33 | 15 | 47 | n | y | y |
| 4 | 31 | 39 | 40 | n | y | n |
| 5 | 49 | 61 | **76** | y | y | y |
| 6 | 82 | 81 | 70 | n | y | n |
| 1, 2 | 88 | 89 | 33 | y | y | y |
| 1, 3 | 88 | 89 | 33 | y | y | y |
| 1, 4 | 77 | 89 | 49 | n | y | n |
| 1, 5 | 70 | 75 | 64 | y | y | y |
| 1, 6 | 83 | 89 | 70 | n | y | n |
| 1, 2, 3 | 88 | 89 | 33 | y | y | y |
| 1, 4, 6 | 83 | 89 | 63 | n | y | n |
| 1, 5, 6 | 83 | 75 | 57 | n | y | y |
| 1, 4, 5, 6 | 83 | 75 | 57 | n | y | y |
| 1, 2, 3, 4, 5, 6 | 83 | 75 | 57 | n | y | y |

ARAT: Action Research Arm Test. BBT: Box and Block Test. NHPT: Nine Hole Peg Test. VPIT: Virtual Peg Insertion Test.

The decision tree models that performed best (i.e., models with maximum balanced accuracy that also correctly predicted the unexpected non-responder) predicted changes in ARAT and BBT with a cross-validated balanced accuracy of 88% and 89%, respectively, and relied only on patient master data. The best decision tree predicting changes in NHPT relied purely on digital health metrics of sensorimotor impairments, yielding a balanced accuracy of 76%. The best linear regression model achieved a balanced accuracy of 94% for the ARAT (independent variables: conventional scales of activity), 75% for the BBT (patient master data and intervention group), and 64% for the NHPT (conventional scales of body functions). The best k-nearest neighbor models achieved a balanced accuracy of 94% for the ARAT (independent variables: patient master data and intervention type), 80% for the BBT (patient master data and intervention type), and 82% for the NHPT (patient master data, digital health metrics, and conventional scales of activity limitations). The best random forest models achieved a balanced accuracy of 78% for the ARAT (independent variables: patient master data), 89% for the BBT (patient master data), and 89% for the NHPT (patient master data and digital health metrics).

For predicting changes in BBT and NHPT, non-linear machine learning models yielded an improvement in accuracy of up to +14% and +25%, respectively, compared to a linear regression approach. The performance of the linear regression models was equal to the non-linear models for predicting changes in the ARAT. For three out of four machine learning models (all despite random forest) that predicted changes in NHPT based solely on digital health metrics or conventional scales of body functions, the ones relying on digital health metrics had superior predictive accuracy by up to +21%. For the random forest, both feature sets had equal predictive power. For all machine learning models combining either digital health metrics or conventional scales of body functions with other feature sets, the ones including digital health metrics outperformed the ones including conventional assessments of body functions by up to 17%.

## 4 Discussion

The objective of this work was to explore the feasibility of predicting the response of individual pwMS to specific upper limb neurorehabilitation interventions by applying machine learning to clinical data and digital health metrics recorded pre-intervention. For this purpose, patient master data, conventional scales describing body functions and activities, as well as upper limb movement and grip force patterns were recorded in 11 pwMS that received eight weeks of neurorehabilitation. Four commonly applied machine learning models (decision tree, random forest, k-nearest neighbor, linear regression) were trained on six different feature sets and combinations thereof. The models were evaluated based on their ability to correctly predict the presence of changes in activity limitations across the intervention and based on their ability to accurately anticipate outcomes for one subject with strong activity limitations at admission but without significant gains across intervention (i.e., an unexpected non-responder).

In summary, changes in ARAT and BBT could be accurately predicted (up to 89% balanced accuracy) by only relying on patient master data (namely age, sex, MS type and chronicity) with slight improvements when adding knowledge about the intervention type (up to 94% balanced accuracy). Moreover, changes in NHPT could be predicted accurately (up to 89% balanced accuracy), but only when providing the models with information about sensorimotor impairments. Assessing these with digital health metrics as provided by the VPIT led to equal or higher predictive accuracies than relying on conventional assessments.

### Machine learning enables a personalized prediction of rehabilitation outcomes in pwMS

These results successfully demonstrate the feasibility of predicting the response of pwMS to specific neurorehabilitation interventions using machine learning and multi-modal clinical and behavioral data. This work especially makes an important methodological contribution, as it is the first attempt towards a personalized prediction of neurorehabilitation outcomes in pwMS. So far, such approaches were rather employed to predict natural disease progression in pwMS [50, 51, 52, 53, 54, 55]. Previous work in neurorehabilitation of pwMS focused on predicting adherence to telerehabilitation [56] or identifying population-level predictors of therapy outcomes through linear regression [13, 14, 15]. For the latter, the models were commonly evaluated by comparing the amount of variance explained by the model with the overall variance, which only provides population-level information and challenges comparisons across models trained on different dependent variables [57, 58]. The presented methodology expands this work by applying an evaluation criteria (balanced accuracy: average of sensitivity and specificity) that can be directly related to the predictive performance of the model for an individual patient and, thus, has higher clinical relevance. Further, the non-linear machine learning models applied in this work were able to strongly improve (up to 25%) the predictive accuracy compared to linear regression approaches. This indicates that the linear relationship often assumed between predictors and rehabilitation outcomes is oversimplified, and that likely non-linear mechanisms underlie the prediction of therapy outcomes. In addition, this stresses the importance of applying non-linear modeling approaches to enable accurate predictions.

The four machine learning models were selected based on their robustness, as they are known to perform well even on rather small datasets [44]. In addition, these models do not require the optimization of specific input parameters or model architectures, as compared to, for example, more complex neural network-based models [44, 59]. Hence, this promises that other researchers can easily adapt the presented methodology. In addition, it should be emphasized that all models were generated in a data-driven manner. In the presented context, such data-driven approaches are preferable over models that require the manual definition of a mathematical formula (e.g., non-linear mixed effect models), which can introduce bias and require advanced knowledge about expected patterns of recovery that is unfortunately often not available.

Ultimately, the presented methodology has the potential to open new avenues in the prediction of neurorehabilitation outcomes in pwMS and other neurological conditions.

### Clinical applicability and mechanisms underlying the prediction of neurorehabilitation outcomes

Patient master data was sufficient to accurately predict changes in the ARAT and BBT. Given that this information is typically available for every patient undergoing neurorehabilitation, such a model could be easily integrated into daily clinical decision making. Therein, the objective output of the model could complement other, often more subjective, information that are used by healthcare practitioners to define patient-specific therapy programs and set rehabilitation goals [4, 5]. As the proposed models seem to be able to identify non-responders to a restorative neurorehabilitation intervention strategy, this could allow to rather focus, for such individuals, on approaches aiming at learning compensatory strategies in order to improve their spectrum of activities, their quality of life and participation in the community. On the other hand, individuals identified as responders might instead benefit from therapy aimed at the neuroplastic restoration of impaired body functions [60, 11]. While the specific mechanisms underlying the predictive power of patient master data remain unclear, we speculate that these data affect multiple aspects that determine the success of neurorehabilitation for example the biological substrates for neuroplasticity, participation in therapy, and learned non-use [60, 11, 61, 56]. The latter might play an especially important role, given that individuals with higher chronicity often showed higher gains in ARAT and BBT (Table 1). Interestingly, knowledge of the intervention type (i.e., task-oriented therapy at high or moderate intensity, or occupational therapy) only led to slight improvements in the prediction of ARAT and BBT scores and solely for the k-nearest neighbor models. While stronger intensity-dependent effects were found in the clinical analysis of this trial [6], its rather minor impact in the analysis presented here might be explained by the reduced number of datasets being available for this work, with only two of them belonging to the occupational therapy group.

Interestingly, information about sensorimotor impairments was necessary to predict changes in the NHPT. This indicates that, most likely, different mechanisms underlie the observed improvements in ARAT and BBT scores compared to NHPT outcomes. While more advanced analysis and large-scale studies would be necessary to fully unravel these mechanisms, we speculate that changes in the ARAT and BBT are influenced by multiple factors such as hand control, voluntary neural drive, weakness, fatigue, and attentive deficits, whereas changes in the NHPT might mainly reflect the recovery of sensorimotor function needed to perform fine dexterous finger movements. One could carefully speculate that this dexterous hand function might be linked to the integrity of the corticospinal tract, which has been shown to be essential for sensorimotor recovery in other neurological disorders [62, 63] and might also play a role in pwMS [64].

### Digital health metrics outperformed conventional scales for predicting changes in the NHPT

The digital health metrics of sensorimotor impairments extracted from the VPIT consistently outperformed conventional scales of sensorimotor impairments when predicting changes in activity limitations, as measured by the NHPT. Hence, we argue that the proposed digital health metrics allow, for this specific application, a superior evaluation of sensorimotor impairments than conventional scales. This is likely because the former provide continuous fine-graded information on ratio scales that might be beneficial for training the machine learning models, compared to the more coarse ordinal scales of conventional assessments. In addition, it seems that especially the two VPIT metrics describing impaired grip force coordination were able to predict changes in the NHPT (Figure 2), which is in line with its expected sensitivity to improvements in dexterous hand control. Also, the superiority of digital health metrics for predicting rehabilitation outcomes might be explained by none of the conventional assessments being able to provide metrics specifically capturing impaired grip force coordination as done by the VPIT.

When comparing the VPIT to other technology-aided assessments, it becomes apparent that most of them focus more on the evaluation of arm movements with less focus on the hand [21, 22, 23, 25, 26, 27], which seems to be especially important for relating impairments to their functional impact. Overall, the VPIT emerges as a unique tool able to provide digital health metrics, which complement the clinically available information about impaired body functions. In addition, the assessments with the VPIT can be performed within approximately 15 minutes per upper limb, thereby showing high clinical feasibility and being well below the time to perform a battery of conventional tests.

### Limitations

A major limitation of this work is the small sample size included for the training and evaluation of the machine learning models. In addition, given the slight imbalance between number of pwMS with and without considerable changes in activity limitations across the intervention, it might be that the models slightly overfitted to the group with more observations. Hence, it is unlikely that the current models would accurately generalize to the heterogeneous population of all pwMS. Further, it is unclear whether the models would be able to predict the effect of a different type of neurorehabilitation intervention or whether therapy parameters would need to be integrated into the model. As any related study, this work is also limited by the specific conventional scales and digital health metrics that were used to quantify impaired body functions and activity limitations. Therefore, it is unclear whether different trends would be observed when considering other conventional or instrumented assessments.

## 5 Conclusions

This work successfully established the feasibility of an individualized prediction of upper limb neurorehabilitation outcomes in pwMS by combining machine learning with multi-modal clinical and behavioral data collected before a neurorehabilitation intervention. Given that non-linear models showed superior predictive power over linear models, this work suggests that the commonly assumed linearity of neurorehabilitation predictors is oversimplified. Lastly, information about sensorimotor impairments was necessary to predict changes in fine dexterous hand control. In these cases, conventional scales of impaired body functions were outperformed in terms of predictive power by digital health metrics, thereby underlining their potential to provide a more sensitive and finegrained assessment.

Future work should focus on validating these results in large-scale populations in order to build models that are more representative of the heterogeneous population of pwMS and can be seamlessly integrated into daily clinical routine. These models should include more holistic information on each individual, including for example information about their psychological status and intrinsic motivation, thereby promising higher prediction accuracies. Also, pivoting from the proposed classification (binary output) towards a regression (continuous output) approach will allow providing a higher level of granularity in the predicted outcomes. Lastly, the inclusion of additional therapy parameters in the models could enable in silico clinical trials, thereby allowing to predict the effects of different therapies for each individual and support a more optimal and data-driven clinical decision making process.

## Acknowledgements

The authors would like to thank Nadine Wehrle for her assistance in data analysis, Monika Zbytniewska for her critical feedback on the manuscript, and Stefan Schneller for contributing to graphical design. This project received funding from the European Union’s Horizon 2020 research and innovation programme under grant agreement No. 688857 (SoftPro) and from the Swiss State Secretariat for Education, Research and Innovation (15.0283-1).

## Ethical approvals

The study was registered at clinicaltrials.gov (NCT02688231) and approved by the responsible Ethical Committees (University of Leuven, Hasselt University, and Mariaziekenhuis Noord-Limburg).

## Conicts of interest

The authors declare no conflicts of interest.

## Data availability

The data presented in this manuscript are available upon reasonable request and under consideration of the ethical regulations.

## Supplementary material

**Table SM3:**
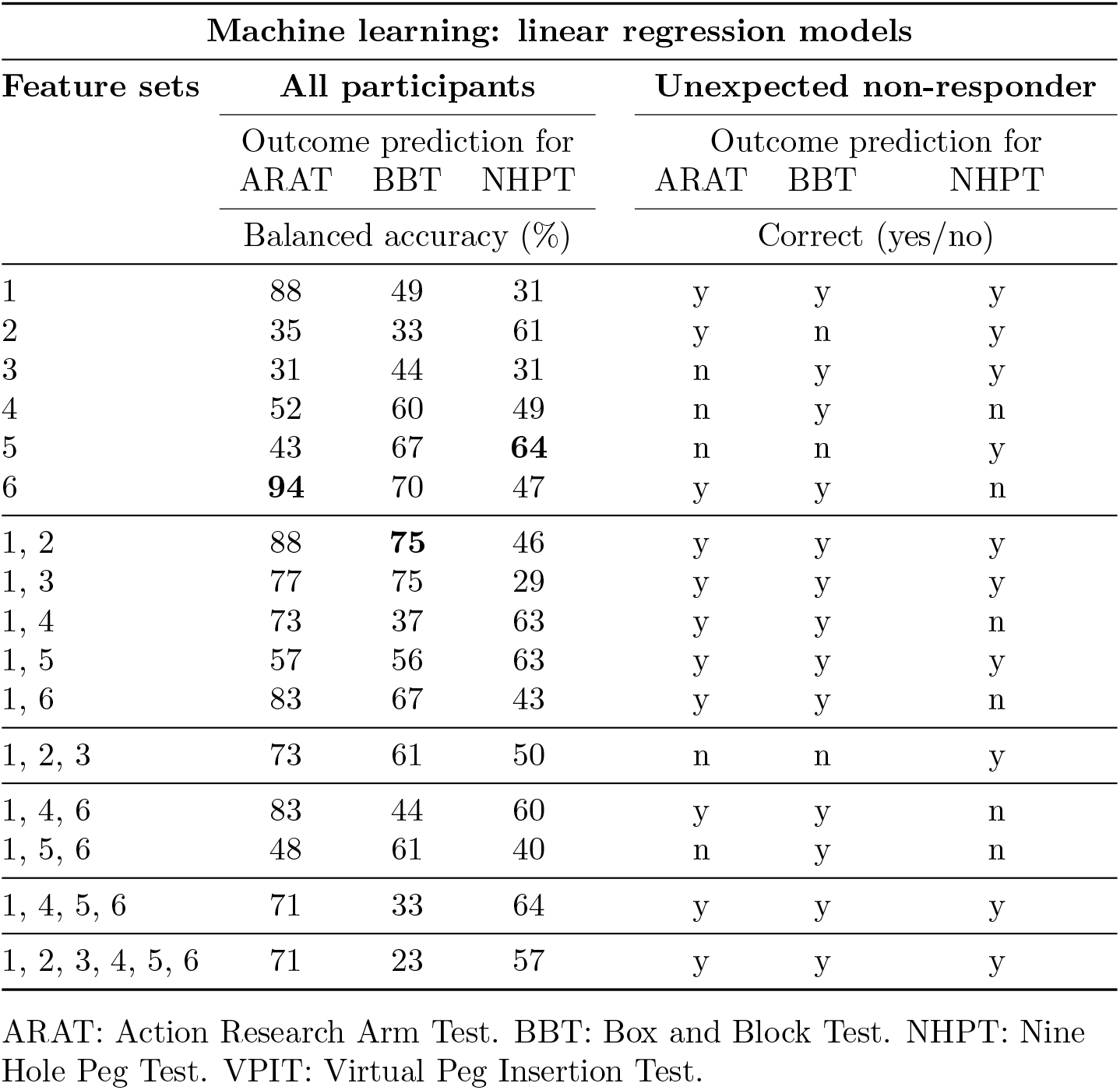
Predicting intervention outcomes using data collected preintervention and a linear regression model. Multiple machine learning models were trained using different feature sets (independent variables, 1-6). The training label indicated whether a considerable change across intervention was observed in a specific conventional score (dependent variable; ARAT, BBT, or NHPT). The models were evaluated in a leave-one-out cross-validation and specifically tested for one individual with strong activity limitations who did not show improvements across neurorehabilitation (referred to as unexpected non-responder). Feature set nomenclature: 1: patient master data (ms type, chronicity, age, sex). 2: intervention group. 3: disability (EDSS, disability group). 4: Conventional scales of body functions (motricity index, static fatigue index, monofilament index, symbol digit modality test, Fahn’s tremor rating scale). 5: Digital health metrics of sensorimotor impairments (ten VPIT metrics). 6: Conv. scale of activity (ARAT, NHPT, BBT). The best performing (accuracy and unexpected non-responder) models relying on the least amount of features are highlighted in bold for each conventional scale.

**Table SM4:** Predicting intervention outcomes using data collected preintervention and a k-nearest neighbor model. Multiple machine learning models were trained using different feature sets (independent variables, 1-6). The training label indicated whether a considerable change across intervention was observed in a specific conventional score (dependent variable; ARAT, BBT, or NHPT). The models were evaluated in a leave-one-out cross-validation and specifically tested for one individual with strong activity limitations who did not show improvements across neurorehabilitation (referred to as unexpected non-responder). Feature set nomenclature: 1: patient master data (ms type, chronicity, age, sex). 2: intervention group. 3: disability (EDSS, disability group). 4: Conventional scales of body functions (motricity index, static fatigue index, monofilament index, symbol digit modality test, Fahn’s tremor rating scale). 5: Digital health metrics of sensorimotor impairments (ten VPIT metrics). 6: Conv. scale of activity (ARAT, NHPT, BBT). The best performing (accuracy and unexpected non-responder) models relying on the least amount of features are highlighted in bold for each conventional scale.

| Machine learning: k-nearest neighbor model |  |  |  |  |  |  |
| --- | --- | --- | --- | --- | --- | --- |
| Feature sets | All participants |  |  | Unexpected non-responder |  |  |
|  | Outcome prediction for |  |  | Outcome prediction for |  |  |
|  | ARAT | BBT | NHPT | ARAT | BBT | NHPT |
|  | Balanced accuracy (%) |  |  | Correct (yes/no) |  |  |
| 1 | 61 | 65 | 33 | y | y | y |
| 2 | 49 | 16 | 53 | n | n | y |
| 3 | 52 | 35 | 33 | y | y | y |
| 4 | 33 | 54 | 40 | n | n | y |
| 5 | 67 | 67 | 70 | y | y | y |
| 6 | 83 | 65 | 64 | n | y | n |
| 1, 2 | **94** | **80** | 33 | y | y | y |
| 1, 3 | 63 | 25 | 35 | y | y | y |
| 1, 4 | 61 | 43 | 72 | y | y | y |
| 1, 5 | 63 | 55 | 70 | y | y | y |
| 1, 6 | 88 | 59 | 35 | n | y | n |
| 1, 2, 3 | 78 | 35 | 35 | y | y | y |
| 1, 4, 6 | 70 | 38 | 47 | n | y | n |
| 1, 5, 6 | 82 | 55 | **82** | n | y | y |
| 1, 4, 5, 6 | 60 | 70 | 60 | n | y | n |
| 1, 2, 3, 4, 5, 6 | 64 | 70 | 52 | n | y | y |

ARAT: Action Research Arm Test. BBT: Box and Block Test. NHPT: Nine Hole Peg Test. VPIT: Virtual Peg Insertion Test.

**Table SM5:**
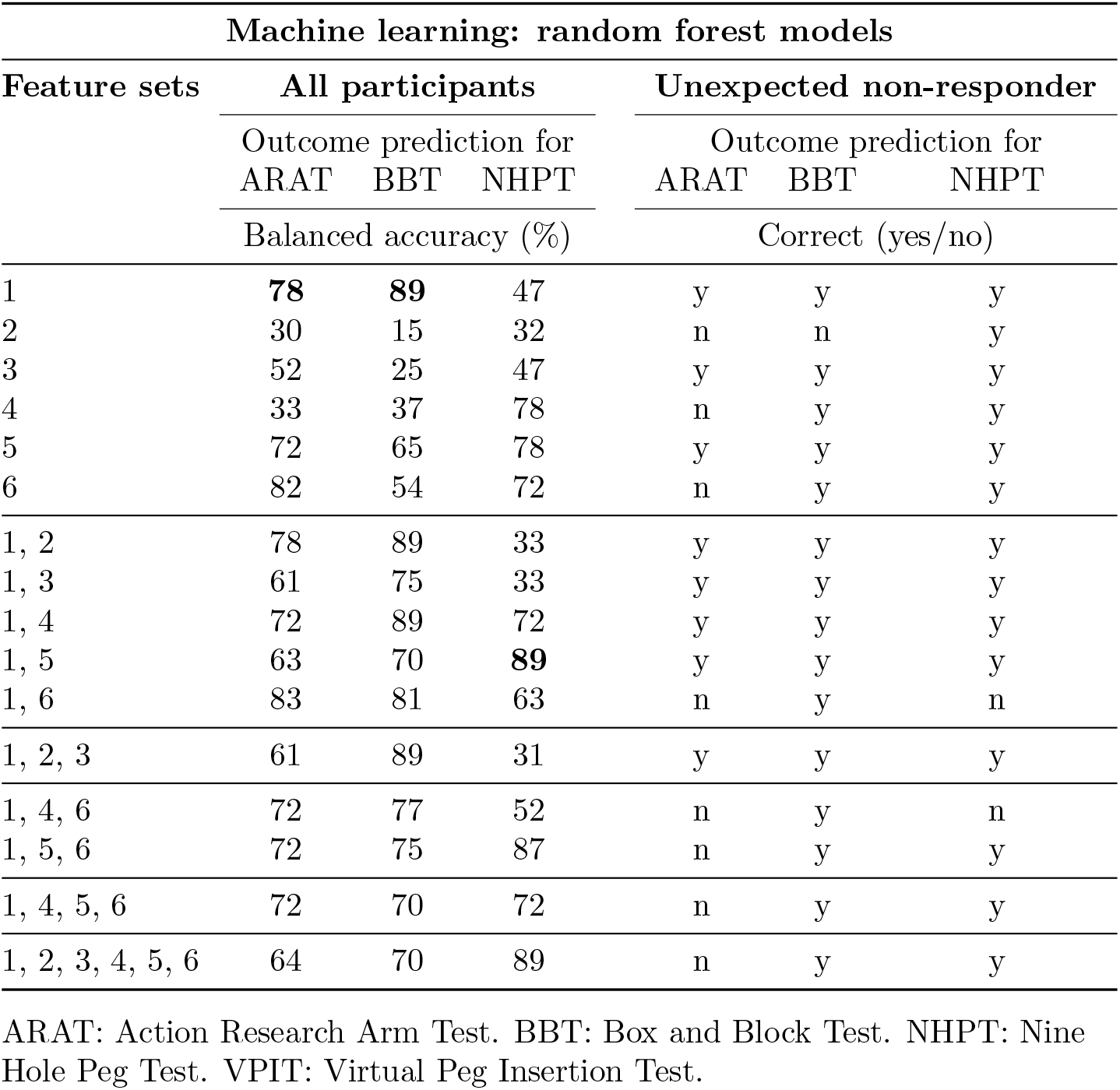
Predicting intervention outcomes using data collected preintervention and random forest models. Multiple machine learning models were trained using different feature sets (independent variables, 1-6). The training label indicated whether a considerable change across intervention was observed in a specific conventional score (dependent variable; ARAT, BBT, or NHPT). The models were evaluated in a leave-one-out cross-validation and specifically tested for one individual with strong activity limitations who did not show improvements across neurorehabilitation (referred to as unexpected non-responder). Feature set nomenclature: 1: patient master data (ms type, chronicity, age, sex). 2: intervention group. 3: disability (EDSS, disability group). 4: Conventional scales of body functions (motricity index, static fatigue index, monofilament index, symbol digit modality test, Fahn’s tremor rating scale). 5: Digital health metrics of sensorimotor impairments (ten VPIT metrics). 6: Conv. scale of activity (ARAT, NHPT, BBT). The best performing (accuracy and unexpected non-responder) models relying on the least amount of features are highlighted in bold for each conventional scale.

